# Coumarin-based blue excitable probe for functional screening of dehalogenase activity

**DOI:** 10.64898/2026.10.02.756220

**Authors:** Sonia Mion, Marnie Mallet, Yoann Louis, Stéphane Fontanay, Florence Ruaudel, Fabiola Bastian, Maïwenn Jacolot, Florence Popowycz, Philippe M. Oger

## Abstract

In the enzymatic toolbox of depollution, hydrolytic dehalogenases are of particular interest for the bioremediation of organohalogenated compounds. To better identify and characterize these enzymes, it is important to develop appropriate sensing methods. Fluorescent probes for dehalogenase detection have been developped in the past years, however, to this date none were available for application using the common FITC parameters (*i.e.* excitation laser at 488 nm). To adress this limitation, we herein report a new fluorescent probe, “COU4-Cl” excitable in the violet (405 nm) and blue (488 nm) wavelengths adapted for the detection and characterization of haloalkane dehalogenases. Its applicability is discussed and evaluated in comparison to the previously described COU2-Cl probe only excitable in the violet (405 nm). This study shows the advantages and complementarity of the two probes, and further demonstrate their application for the discrimination of dehalogenase activity in whole cell assays.

**GRAPHICAL ABSTRACT:** 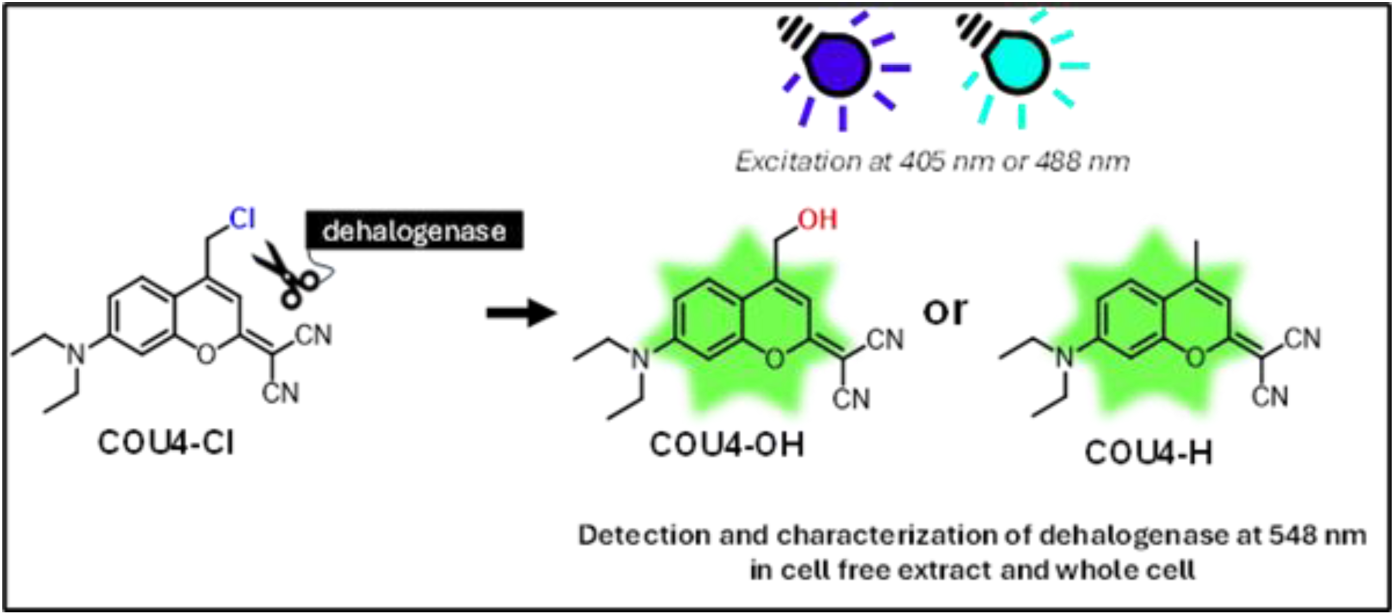

## INTRODUCTION

Many organohalogenated compounds (OHCs) have been synthetized and used since the last century with applications in agriculture, pharmaceutical and material sciences.^1^ However, their chemical stability results in environmental persistence and increasing ecological risks, raising concerns about ecological safety and human health.^1^ The persistence of these compounds is largely due to the strength of the carbon−halogen bond. The thermodynamic stability of these bonds increases significantly from C−Br (318.0 ± 8.4 kJ.mol^-1^) to C−Cl (394.9 ± 13.4 kJ.mol^-1^), culminating in the high strength of C−F bonds (513.8 ± 10.0 kJ.mol^-1^).^2^ Biodegradation of OHCs by microbial enzymes offers an efficient and ecofriendly route to detoxify and recycle these compounds. Among these enzymes, dehalogenases play a central role by catalyzing the cleavage of carbon−halogen bonds in diverse halogenated substrates, including problematic OHCs.^3,4^ By removing halogen substituents, these enzymes detoxify halogenated compounds and convert them into H or OH derivatives (by reductive or oxidative enzymatic activity respectively), making them more prone to further microbial degradation.^5^

Considering their potential for OHCs bioremediation, it is crucial to optimize the detection and characterization of new environmental and engineered dehalogenases. For many years, two methods mainly have been used for the functional detection and characterization of such enzymes: the use of colorimetric pH indicators such as phenol red or bromothymol blue based on the work of Holloway and co.^6^ and the colorimetric detection of halide, based on the work of Iwasaki and co.^7^ These methods are limited in terms of sensitivity and can involve the manipulation of toxic substances such as the mercuric thiocyanate.^7^

More recent alternative approaches to these methods have been developed, relying for instance on the detection of fluorescent signals using fluorescent pH detectors.^8,9^ In comparison to colorimetric assays, fluorescence detections can offer many advantages such as higher sensitivity, lower limits of detections, lower susceptibility to interferences and can be used in different applications such as fluorescence-activated cell or droplet sorting (FACS and FADS) screening, microscopy experiments or *in situ* detection methods.

The design and optimization of fluorogenic substrates for haloalkane dehalogenases has started to build since the work of Dockalova and co. describing the first fluorogenic substrates for haloalkane dehalogenases^10^, **Scheme 1**. Based on a coumarin for **COU-Br** or BODIPY structure for **BDP-Cl**, the dehalogenation of these molecules induced the formation of a hydroxylated product with a higher fluorescence emission intensity than the substrate, allowing the detection and characterization of dehalogenase enzymes based on fluorescence and the in-depth study of their catalytic mechanism. They further developed new coumarin probes **COU-1**, **COU-2** and **COU-3** as described in a thesis report^11^ and a recent paper^12^ where **COU-3** was successfully applied in a high throughput FADS system to screen an engineered dehalogenase LinB mutants library. Among these substrates, **COU-Br**, **COU-2** and **COU-3** are detectable using violet excitation (405 nm), and only **COU-1** and **BDP-Cl** can be detected using the more common FITC parameters (488 nm). Due to the use of differentially substituted coumarin derivatives in this work, we will use a specific notation highlighting the substituting group present on the coumarin, *i.e.* **COU-2** which present a Cl substitution will be noted **COU2-Cl** throughout this paper, while **COU2-H** and **COU2-OH** will refer to the hydrogenated and hydroxylated derivatives). The stock shift of **BDP-Cl** and its hydroxylated products **BDP-OH** is quite small (16 nm between absorption and emission maxima for **BDP-Cl**, 18 nm for **BDP-OH**), and **COU-1** presented a high level of spontaneous hydrolysis in phosphate buffer (4% conversion in 24h) and a low solubility. These results called for further development of new fluorescent probes usable in FITC parameters.

**Scheme 1.**
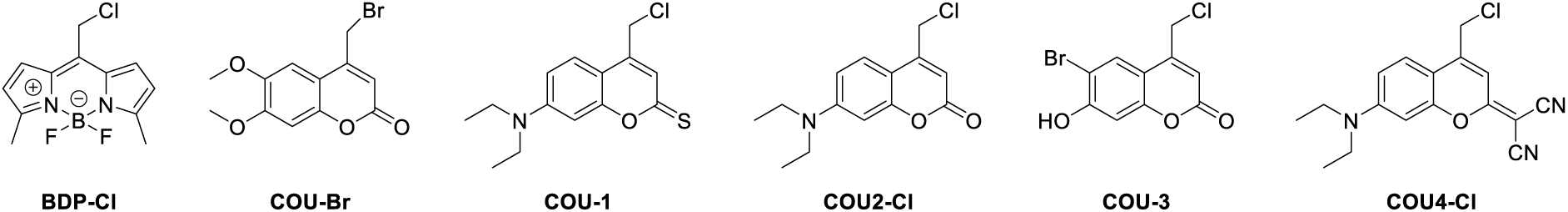
Molecular structures of BODIPY (BDP) and coumarin (COU) probes previously designed to detect dehalogenase activity: **BDP-Cl**, **COU-Br**, **COU-1**, **COU2-Cl**, **COU-3** described in precedent papers^10,12^ and the newly synthetized **COU4-Cl** described in this study.

Among the previously described fluorescent substrates, **COU2-Cl** is a molecule of great interest with regards to its stability, solubility and ability to detect and characterize the activity of the haloalkane dehalogenase DmmA.^12^ However, this probe can only be excited in the violet and UV range (λ*^exc^*_max_ at 389 nm) making it unsuitable for many applications and instruments, which will often possess a laser at 488 nm. To circumvent this issue, we synthesized a new probe **COU4-Cl**, inspired by the structure of **COU2-Cl** with the introduction of a cyano group to shift the molecule’s fluorescence spectrum, thereby enabling enzyme detection using the common FITC parameters.^13^ This work presents the characterization of **COU4-Cl** and its comparison with **COU2-Cl** for the detection and evaluation of dehalogenase enzyme in cell-free extracts and in whole bacteria cells.

## RESULTS AND DISCUSSION

### Characterization of the new COU4-Cl fluorescent substrate in comparison with COU2-Cl

**COU4-Cl** was designed with the objective of creating a molecular probe that could be detected using a 488 nm excitation laser. As the coumarin probe **COU2-Cl** had already presented good capacity (solubility, stability and sensibility) for dehalogenation detection by fluorescence, the 7- diethylamino-coumarin structure was kept.^11,12^ It was previously shown that the incorporation of two cyano groups at position 2 led to dicyanomethylenecoumarinylmethyl derivatives with maximal absorption around 500 nm in the 7-diethylamino series such as **COU4-OH**^14^. Based on these data the **COU4-Cl** molecule was designed and synthetized. In order to characterize the new probe in comparison with its putative dehalogenated derivatives **COU4-H** and **COU4-OH** resulting from a reductive or oxydative dehalogenation, as well as with the described**COU2-Cl** probe, we chemically synthesized the two halogenated probes (COU2-Cl and COU4-Cl) and their degradation substrates (**COU2-H**, **COU2-OH** and **COU4-H, COU4-OH**) (**Figure 1.A, C**). The photochemical parameters were assayed in phosphate buffer at pH 8 with 10% (v/v) of DMSO. The use of co-solvent was necessary due to the limited solubility of the fluorescent probes. DMSO was chosen as it has already been used to characterize dehalogenase fluorescent probes^10^ and is known to be well tolerated by haloalkane dehalogenases.^15^ It was not used at more than 10% to limit the impact on dehalogenase activity.

**Figure 1.**
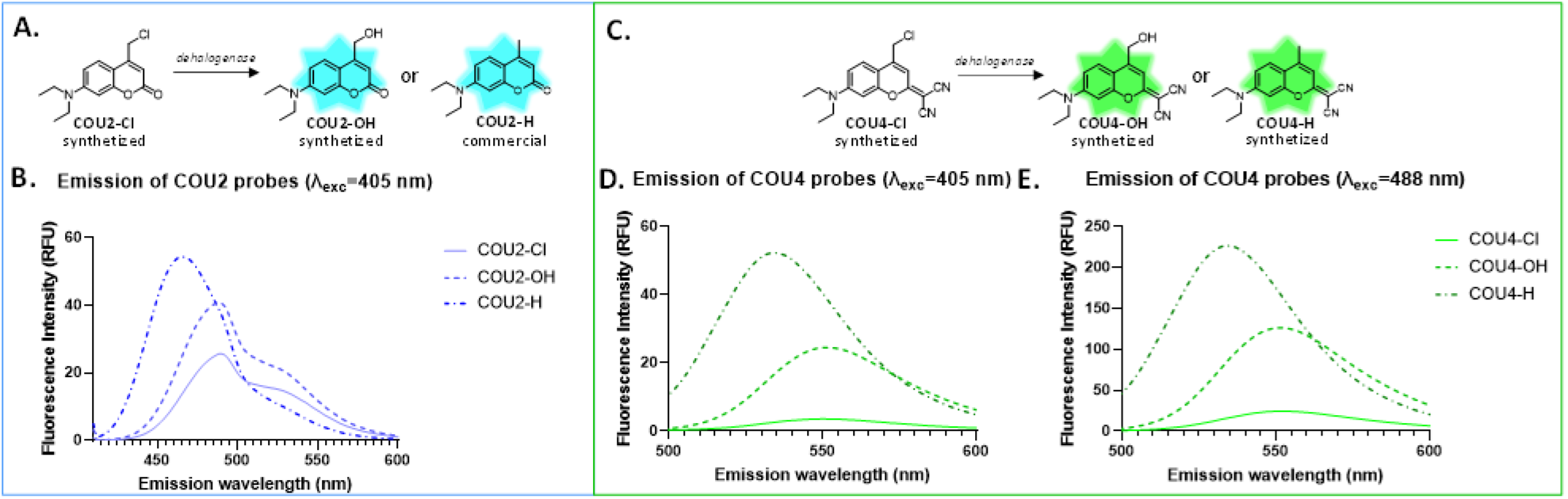
Schematic representation of the probes **COU2** (A) and **COU4** (C) and their potential dehalogenation products and comparison of fluorescence intensity of emission spectra of **COU2** probes after excitation at 405 nm (B) and of **COU4** probes after excitation at 405 nm (D) or at 488 nm (E). All spectra measured with 10 µM of probe in 50 mM phosphate buffer/DMSO mixture (90:10, v/v), pH 8.0.

Looking at the emission/excitation spectra, **COU2-Cl** was excitable in the UV range (389 nm), with a maximum emission in the cyan-blue range (496 nm) in coherence with the literature^11,12^, whereas the desired shift toward higher excitation wavelength was successfully obtained for **COU4-Cl** (**Table 1**, Erreur ! Source du renvoi introuvable., Erreur ! Source du renvoi introuvable.). Interestingly, **COU4-Cl** can still be excited in the UV range (**Figure 1.D**) but has a maximum excitation in the cyan range (480 nm), and can be detected in the green range (548 nm) (**Table 1**). As observed for the **COU2** probe, the hydrolysed form **COU4-OH** presents similar excitation/fluorescence spectra than **COU4-Cl**, and a small shift toward smaller wavelength (530 nm) is observed for the reduced form **COU4-H** (**Table 1**, Erreur ! Source du renvoi introuvable., Erreur ! Source du renvoi introuvable.).). More importantly, the dehalogenated products **COU4- OH** and **COU4-H** are significantly more fluorescent than the chlorinated form **COU4-Cl** after excitation at 405 nm or 488nm (**Figure 1**). The quantum yield of the product forms (**COU4-OH** and **COU4-H**) are respectively 1.5 to 2 times higher than the substrate form **COU4-Cl** (**Table 1**). The absolute quantum yield of the **COU4** family which varies between 18.8 % for **COU4-Cl** and 41.97 % for **COU4-OH** is higher than the **COU2** family which varies only between 4.35 % for **COU2-Cl** and 7.32 % for **COU2-H** (**Table 1**), and thus **COU4** presents a better fluorescence potential. Compared to **COU2**, the dehalogenation could therefore be detected by an augmentation of fluorescence with more sensitivity, using one or both of the common 405 nm (violet) and 488 nm (blue) excitation lasers available in several screening instruments such as microplate readers, fluorescence microscopes or cytometers, whereas **COU2** can only be detected using the 405 nm excitation laser (**Figure 1**).

**Table 1.** Photophysical properties of fluorescent substrate.

| | $\lambda_{\max}^{\text{exc}}$(nm) | $\lambda_{\max}^{\text{fl}}$(nm) | $\Phi_{\text{fl}}$(%) |
| --- | --- | --- | --- |
| **COU2-Cl** | 389 | 496 | $4.35 \pm 0.43$ |
| **COU2-OH** | 389 | 496 | $7.23 \pm 0.04$ |
| **COU2-H** | 385 | 460 | $7.32 \pm 0.53$ |
| **COU4-Cl** | 483 | 548 | $18.80 \pm 1.31$ |
| **COU4-OH** | 483 | 548 | $41.97 \pm 0.65$ |
| **COU4-H** | 483 | 530 | $28.10 \pm 1.40$ |

Absorption ( $\lambda_{\max}^{\text{exc}}$ ) and emission ( $\lambda_{\max}^{\text{fl}}$ ) maxima and quantum yields ( $\Phi_{\text{fl}}$ ) of each coumarin compound measured in solutions of 50 mM phosphate buffer/DMSO mixture (90:10, v/v), pH 8.0, c = 10 $\mu\text{M}$ for $\lambda_{\max}^{\text{exc}}$ and $\lambda_{\max}^{\text{fl}}$ and 7 $\mu\text{M}$ for $\Phi_{\text{fl}}$ .

### Application of COU4-Cl for the detection and characterization of dehalogenase activities

#### COU4-Cl and COU2-Cl can be used to screen specific activity of diverse haloalkane dehalogenase

**COU2-Cl** has previously been used to detect and characterize the activity of the haloalkane dehalogenase DmmA isolated from a marine microbial consortium.^11,12,16^ In order to demonstrate the potential of **COU4-Cl** to detect and characterize enzymatic activities and compare it with **COU2-Cl**’s, the activities of 15 diverse commercial wild-type haloalkane dehalogenases (HLDs) were assayed by monitoring the fluorescence increase corresponding to the formation of the hydrolyzed product **COU4-OH** or **COU2-OH** respectively (**Figure 2**, Erreur ! Source du renvoi introuvable.), in addition the consumption of the chlorinated product and formation of hydroxylated product was confirmed for LinB and DmmA dehalogenase using HPLC-FLD (Erreur ! Source du renvoi introuvable.).

**Figure 2.**
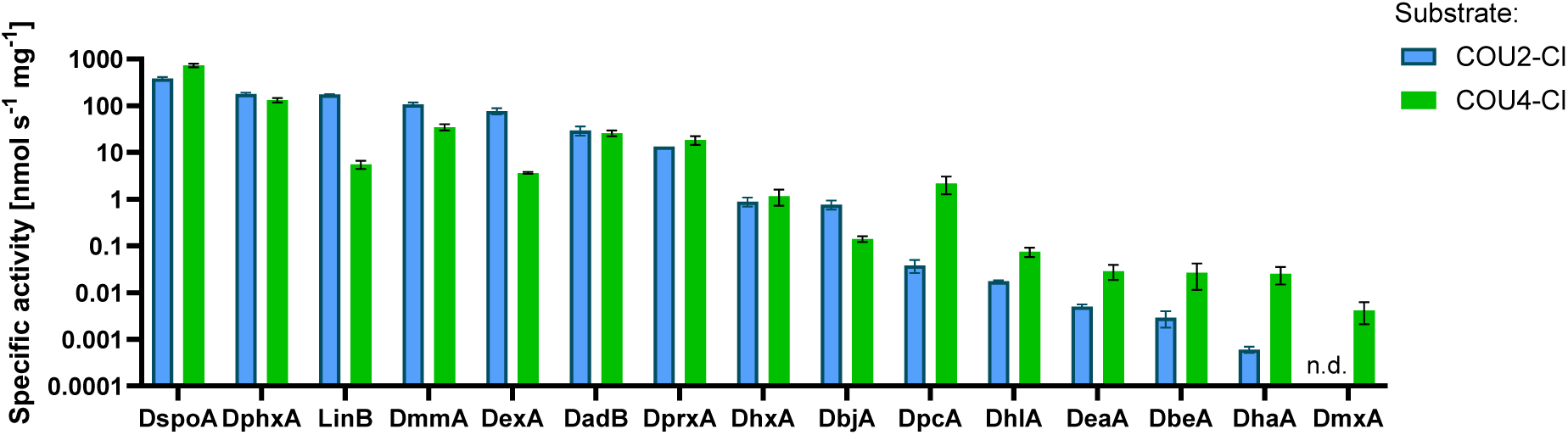
Specific activities of wild types HLDs on **COU4-Cl** and **COU2-Cl**. Specific activities with standard errors for HLDs on **COU2-Cl** (blue with border) and **COU4-Cl** (green, no border) recorded by fluorescent measurements of **COU2-OH** (λ_exc_ = 389 nm, λ_em_ = 496 nm) or **COU4-OH** (λ_exc_ = 484 nm, λ_em_ = 548 nm) product apparition respectively. n = 3 replicates. n.d. = not detected

Interestingly, even though the coumarin substrates are quite bulky compared to most common alkane substrates such as 1-chlorobutane, it was shown that on **COU4-Cl,** activities were detected for all the tested enzymes, whereas on **COU2-Cl,** activities were detected for 14 of the 15 HLDs, overall showing the potential of the two probes to detect activities as small as 0.61 pmol.s^-1^.mg^-1^ (DhaA on **COU2-Cl**) and 4.15 pmol.s^-1^.mg^-1^ (DmxA on **COU4-Cl**) and up to 382 nmol.s^-1^.mg^-1^ (DspoA on **COU4-Cl**) and 728 nmol.s^-1^.mg^-1^ (DspoA on **COU4-Cl**). It should be noted than even if several enzymes presented the same order of activity toward the two probes, DpcA, DhlA, DeaA, DbeA, DhaA showed a higher specific activity for **COU4-Cl** than **COU2-Cl**, and no activity was detected for DmxA on **COU2-Cl** whereas the enzyme presented a detectable activity of 4.15 pmol.s^-1^.mg^-1^ toward **COU4-Cl**. **COU4-Cl** therefore appears as a more sensitive dehalogenase detection probe with a broader range of dehalogenases detected than **COU2-Cl**.

### COU2-Cl and COU4-Cl can be used to characterize steady state kinetics of diverse haloalkane dehalogenase

**COU4-Cl** and **COU2-Cl** were further used to determine the steady-state kinetic parameters of the most active enzymes (DspoA, DphxA, LinB, DmmA, DexA, DadB, and DprxA) toward these substrates using the fluorescence conversion curves recorded at different substrate concentration.

In general, the affinities of the seven tested enzymes was greater for **COU2-Cl** than for **COU4- Cl** (**Table 2**) and higher than the affinities described for smaller substrates such as 1,2- dibromoethane or 1,3-dibromopropane.^10,17,18^ DmmA and LinB presented the highest affinities toward **COU2-Cl** (K_m_ of 4.95 µM and 4.59 µM respectively) and DmmA and DphxA from *Phaeobacter* sp. CECT 5382^17^ the highest affinities toward **COU4-Cl** (K_m_ of 12.08 µM and 13.09 µM respectively). This greater affinity for bulky substrates had already been described for DmmA and LinB from *Sphingobium japonicum* UT26^19^ toward COU-Br and BDP-Cl and **COU2-Cl**^10,12^, reenforcing the hypothesis of being closer to the natural substrates of these enzymes.

**Table 2.**
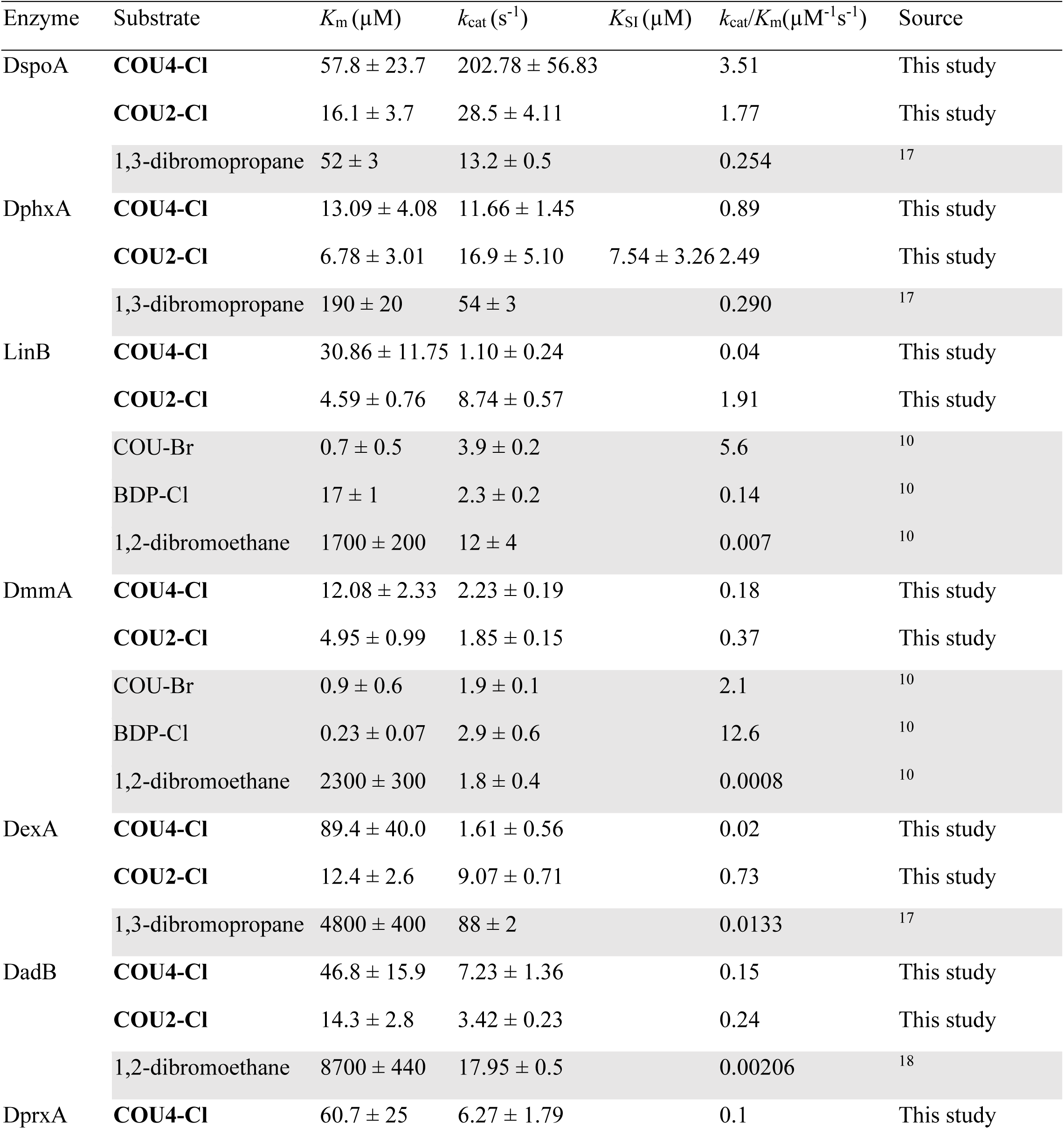
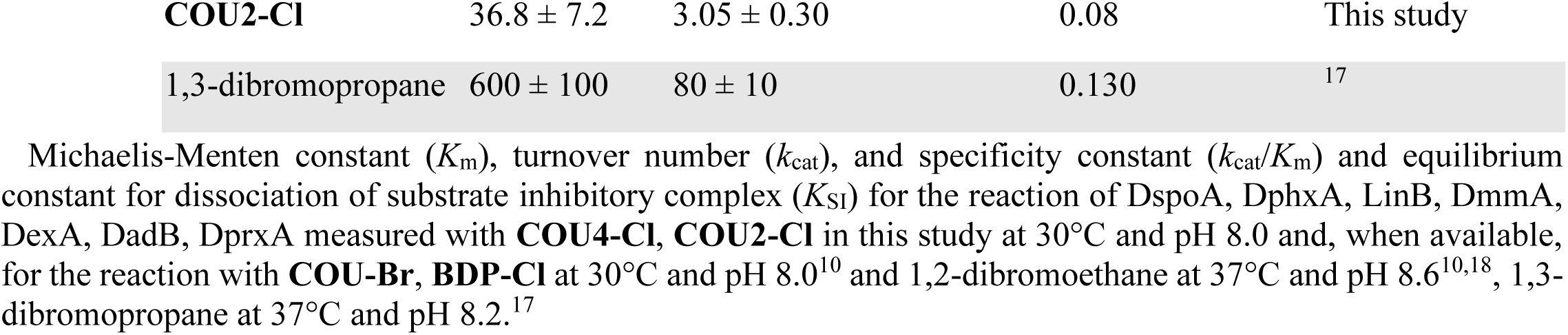
Steady-state kinetic parameters.

Among the seven HLDs, DspoA showed the highest turnover number k_cat_ toward the two fluorescent substrates, reaching a rare level of 202.78 s^-1^ toward **COU4-Cl** and 28.5 s^-1^ toward **COU2-Cl**. These elevated values translate in the highest specificity constant k_cat_/K_m_ observed toward **COU4-Cl** (3.51 µM^-1^.s^-1^), and one of the three highest k_cat_/K_m_ observed toward **COU2-Cl** (1.77 µM^-1^.s^-1^), in the same range as DphxA (2.49 µM^-1^.s^-1^) and LinB (1.91 µM^-1^.s^-1^). Interestingly, DspoA has only been identified recently by advanced database mining, showing a good specific activity toward 1,3-dibromopropane and a high temperature stability (with an apparent melting temperature of 60°C).^17^ In regard of this new results, this enzyme appears as a new interesting tool for the dehalogenation of a broad range of substrates, from small brominated substrates to bulky chlorinated ones.

Substrate inhibition was observed in the reaction of DphxA with **COU2-Cl**. The kinetic analysis provided equilibrium constant for dissociation of substrate inhibitory complex K_SI_ = 7.54 ± 3.26 µM. It is not surprising as substrate inhibition is the most common deviation from Michaelis– Menten kinetics, occurring in approximately 25% of known enzymes. It can be attributed to the formation of an unproductive enzyme–substrate complex after the simultaneous binding of two or more substrate molecules to the active site or by substrate binding to the enzyme–product complex.^20^ Here, the phenomenon is observed only for this enzyme, and was not observed with **COU4-Cl** in these conditions. It has already been described for HLDs and bulky substrates such as in the reaction of DmmA with BDP-Cl^10^, **COU1-Cl** and **COU3-Cl.**^12^

**COU4-Cl** is a useful probe for the characterization of haloalkane dehalogenase activity. Even with a chemical structure highly similar to **COU2-Cl**, difference in activities and kinetic parameters are observed for several enzymes towards the two probes. These different behaviours (substrate inhibition, higher specificity) toward substrates which are very similar in structure highlight the interest of using the two probes in combination for further investigation of haloalkane dehalogenase catalytic mechanisms.

#### COU2-Cl and COU4-Cl can be used in whole cell assay for dehalogenase screening

**COU4-Cl** has been developed as a new tool for the screening and detection of dehalogenase activity. After validating that the probe was adapted for the detection and characterization of enzymes in the form of cell free extracts, **COU4-Cl** was compared to **COU2-Cl** for its capacity to detect dehalogenase activity in a whole-cell assay. E. coli expressing the dehalogenase LinB, and empty vector control cells were incubated with the probes and the fluorescence was followed over time. In these conditions, after only 30 minutes the fluorescence was significantly different between the control and the LinB-expressing cells presenting a fluorescence 18 times and 4 times higher in dehalogenase expressing cells than in empty vector control with **COU2-Cl** and **COU4-Cl** respectively (**Figure 3**, Erreur ! Source du renvoi introuvable.). The better detection using **COU2- Cl** is certainly due to the higher activity of LinB towards **COU2-Cl** than **COU4-Cl** (**Figure 2**). Nevertheless, these results show that **COU4-Cl** can also be applied to efficiently screen whole-cell bacteria for dehalogenase activity. This proof of concept paves the way for the application of the probe for high throughput screening of engineered enzyme libraries, metagenomic libraries or environmental bacteria, using 96 well plate or 384 well plate reader. The fluorescence activity could also be used to develop fluorescent activated cell sorting (FACS) screening as it was recently demonstrated with the probe **COU3-Cl**^12^; **COU4-Cl** presenting the advantages of being usable with two laser excitation (405 nm or 488 nm). For this special application requiring the encapsulation of cell and substrate in water-in-oil droplet, special attention should be made on the lipophilic behaviour of **COU4-Cl**. Indeed, **COU2-Cl** has already been proven unsuitable for this application as it is leaking rapidly from aqueous to oil phase.^12^ In the case where **COU4-Cl** should present the same default, further functionalization of the coumarin structure will need to be investigated to improve the retention of the probe in the aqueous phase.^21^

**Figure 3.**
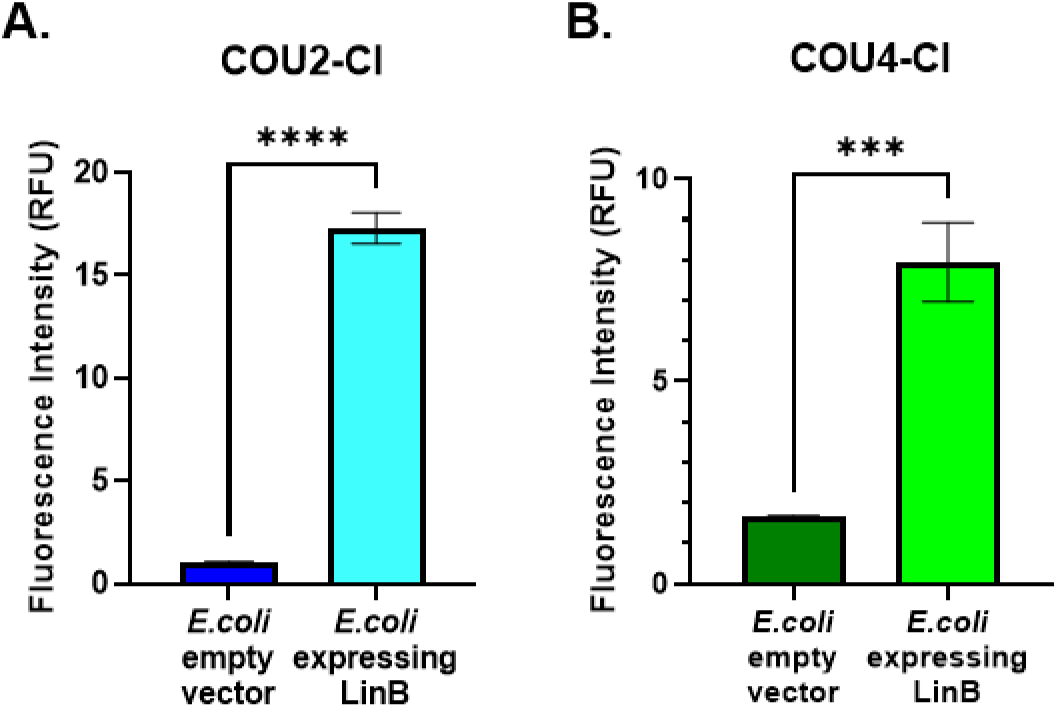
Fluorescent detection of *E. coli* expressing dehalogenase. Fluorescence intensity after 30 min incubation at 30°C of *E. coli* cells with 8 µM **COU2-Cl** (A) or **COU4-Cl** (B), 10 % (v/v) DMSO in phosphate buffer at pH 8. n = 3 replicates for each condition. Fluorescence intensity measured in RFU, λ_exc(A)_ = 389 nm, λ_em(A)_ = 496 nm, λ_exc(B)_ = 483 nm, λ_em(B)_ = 548 nm. **** p- value <0.0001, *** p-value <0.001 of unpaired t-test.

## CONCLUSION

Here we describe a novel coumarin probe **COU4-Cl** specifically designed for the detection and characterization of haloalkane dehalogenase activities, and adapted for whole-cell screening in common fluorescent instruments, and thereby enlarging the toolbox available for dehalogenase detection. In comparison with the known **COU2-Cl** it can be used with both excitation’s wavelength of 405 and 488 nm for a wider range of optical devices, and was shown in this paper to be able to detect the activity of many haloalkane dehalogenases. When used to determine the kinetic parameters of HLDs, the two probes revealed distinct enzyme affinities and catalytic efficiencies, despite their structural similarity. These differences highlight the value of both substrates in tandem to gain deeper insights into the catalytic mechanisms of HLDs. Interestingly, we showed that **COU4-Cl** and **COU2-Cl** could be used to detect reductive dehalogenation as the expected products **COU2-H** and **COU4-H** presented an elevated fluorescence, even if this hypothesis requires further confirmation using reductive dehalogenase on the probes in enzymatic assay. Finally, the two probes were further demonstrated able to detect dehalogenase activities in whole live cells, paving the way to their application for screening environmental and engineered libraries and help tackle OHCs environmental pollution. In perspective, these probes could be used to develop high throughput assays such as FADS following the example of Faldynova and co.^12^, and adapted as iodinated, brominated or even fluorinated probes, to detect new dehalogenases targeting concerning OHCs pollutants such as PFAS.

## EXPERIMENTAL SECTION

### Synthesis of coumarin probes

**COU2-H** is commercially available as 7-(diethylamino)-4-methyl-2*H*-chromen-2-one. The synthesis of **COU2-Cl**, **COU2-OH**, **COU4-OH** and **COU4-H** were already described elsewhere ^13,22–24^ and we applied PCl_5_ chlorination of COU4-OH to obtain the novel probe **COU4-Cl** (*see Supplementary Information for details*).

#### Photophysical Properties

##### Excitation/emission spectra

Excitation and emission spectra were recorded with a spectrofluorometer FP-8300 (Jasco) at 25 °C. The reaction mixture consisted of 90% (v/v) phosphate buffer (pH 8) and 10% (v/v) DMSO with the total concentration of the substrates of 10 µM. Excitation spectra were measured by analyzing the intensity of light emitted at the maximal emission wavelength. Emission spectra were measured by exciting the sample with light corresponding to the maximal excitation wavelength. **Fluorescence quantum yield**

Absolute fluorescence quantum yield was measured in quartz cuvette using a spectrofluorimeter FS5 (Edimburgh Instrument) equipped with an integrating sphere SC-30. 3 mL of each probe was prepared at 7 µM with 10% (v/v) DMSO in phosphate buffer (pH 8), the reference was a solution of phosphate buffer (pH 8) with 10% (v/v) DMSO. The measures were done in triplicate for each probe, at excitation wavelength of 389 nm for **COU2** and 483 nm for **COU4**.

#### Application to enzymatic detection and characterization

##### Specific activity assay and steady-state kinetics

Activity measurements and steady-state kinetic experiments were performed in microtiter plates at 30 °C using a SAFAS Xenius XM (Safas, Monaco) spectrofluorimeter equipped with a xenon flash lamp. The reaction was initiated by addition of 20 µL of substrate solution to a total volume of the reaction mixture of 200 µL. Fluorescence intensity was measured from the top using excitation/emission monochromators set to wavelengths 389/496 nm for **COU2-Cl** and 483/548 nm for **COU4-Cl** with a bandwidth of 10 nm. Before measurement, the microtiter plate was shaken for 3 s and then the increase in fluorescence intensity was monitored at regular time intervals. The reaction took place in in triplicates and using control wells with phosphate buffer without enzyme for monitoring spontaneous hydrolysis of the substrate. The cell free extracts corresponding to the recombinant expression and production in *E. coli* of haloalkane dehalogenase were purchased from Enantis (Brno, Czech Republic) and are described in Erreur ! Source du renvoi introuvable..

##### HPLC-FLD detection and identification of coumarin products

Microtiter plates from detection activity were centrifuged 10 minutes at 3,000 g. Then, 100 μL of supernatant was collected and transferred to a clean microtiter plate and kept at 4 °C in the measuring device (HPLC-UV system Thermofisher Scientific Vanquish). Analysis was carried out using a reverse phase silica Accucore^TM^ C18 (Thermofisher Scientific, diameter 2.1 mm, particule size 2.6 µm, 150 mm length). 5 µL of sample was injected and chromatographic separation was carried out using a mobile phase composed of acetonitrile/water applied in the form of a gradient. Phase started from acetonitrile/water (1:4, v/v), then a linear gradient was applied to reach 100% acetonitrile at 12 minutes of analysis, kept until the end at 20 minutes. The flow rate of the mobile phase was set at 0.3 mL.min^-1^ and the temperature of the column maintained at 30 °C. Fluorometric detection was achieved using the fluorescence detector of the device (Vanquish^TM^ VC-D-50-A, 8 µL), with excitation/emission wavelength of 389/496 nm for COU2 and 483/548 nm for COU4.

##### Recombinant expression of haloalkane dehalogenase LinB

The codon-optimised gene *linB* was commercially synthesized and cloned into recombinant plasmid pET28, the pET28::linBHis plasmid was then transformed into *Escherichia coli* BL21 (DE3) pLysS cells. For overexpression of LinB, 500 µL of precultivated culture was used to inoculate 50 mL of LB media containing kanamycin (50 µg.mL^-1^) and chloramphenicol (12.5 µg.mL^-1^), incubated at 37°C and 115 rpm agitation for 3-4 hours until an OD600 of 0.6 was reached. The culture was cooled at 20°C, and induction of dehalogenase expression was initiated by addition of IPTG to a final concentration of 500 µM. After induction the culture was incubated at 20°C and 115 rpm overnight.

##### Detection of activity using whole cell

1 mL of *Escherichia coli* BL21 (DE3) pLysS pET28::linBHis induced cells were centrifuged at 7000 g for 1 min and washed in 50 mM Phosphate buffer (pH 8.0) two times. Cells are then resuspended in 50 mM Phosphate buffer pH (8.0) at a final OD600 of 2. Activity measurements were performed in microtiter plates at 30 °C using a SAFAS Xenius XM (Safas, Monaco) spectrofluorimeter equipped with a xenon flash lamp. The reaction was initiated by addition of 20 µL of cells solution and 20 µL of substrate solution to a total volume of the reaction mixture of 200 µL (80% (v/v) phosphate buffer (pH 8.0) with 10% (v/v) DMSO, 10% cells solution (v/v) with a final concentration of substrate of 8 µM). Fluorescence intensity was measured from the top using excitation/emission monochromators set to wavelengths 389/496 nm for COU2 and 483/548 nm for COU4 with a bandwidth of 10 nm. Before measurements, the microtiter plate was shaken for 3 s (Amplitude 3, 4Hz).

### Data analysis

In the case of steady-state kinetics with fluorogenic substrates, the observed fluorescent signal was converted into product concentration based on calibration curves constructed from fluorescence intensity values after total conversion of substrate solutions with various concentrations (from 0.8 to 80 µM of substrate). The effects of spontaneous hydrolysis were subtracted from the converted data. Data were analyzed conventionally by measuring initial rates of the reactions proceeding at various substrate concentrations. The rates derived by fitting linear functions to product concentration increase in time were plotted as a function of substrate concentration and fitted by nonlinear regression to Michaelis-Menten kinetic model using GraphPad Prism 10 Software (GraphPad Software Inc.; USA).

## SUPPORTING INFORMATION

Supporting information: additional figures on absorption and emission spectra of coumarin substrates and products, values of specific activities of wild type HLDs toward **COU2-Cl** and **COU4-Cl**, HPLC-FLD chromatograms of **COU2-Cl** and **COU4-Cl** digested by haloalkane dehalogenases, kinetic of dehalogenase activity in whole cell by fluorescence detection using **COU2-Cl** or **COU4-Cl** as substrate and a description of probe **COU4-Cl** synthesis (DOC).

## ACKNOWLEDGMENTS

This work was financially supported by the French Agence Nationale de la Recherche (ANR) in the frame of the project METHALO ANR-22-CHLD-003 to PMO (Post-Doctoral support for SM), the French Ministry of Higher Education, Research and Innovation (Ph.D. Support for MM), the DTAMB Development project 2022 DEMETER to PMO (support for FACS implementation) and the Depollution program from the Office of cross-disciplinary and interdisciplinary initiative (Mission pour les Initiatives Transverses et Interdisciplinarité - MITI CNRS) to PMO. We would like to acknowledge the contribution of the FR BioEEnViS platform DTAMB. The authors wish to thank Dr. Julien Houel (Université Lyon 1, Light-Matter Institute) for his help and advices on quantum yield measurements and Prof. Zbynek Prok from Masaryk University, Czech Republic for the enriching exchanges on fluorescent probe for dehalogenase detection. This work benefited from the remarkable analytical facilities of “Centre Commun de RMN” and “Centre Commun de Spectrométrie de Masse” from the University Lyon 1 and the help of their managing staff.

## Notes

### Competing Interest Statement

The authors have declared no competing interest.

