## Supplementary Info for "Coumarin-based blue excitable probe for functional screening of dehalogenase activity"

### Supporting information

#### 2D and 3D Absorption and emission spectra of coumarin substrates and products

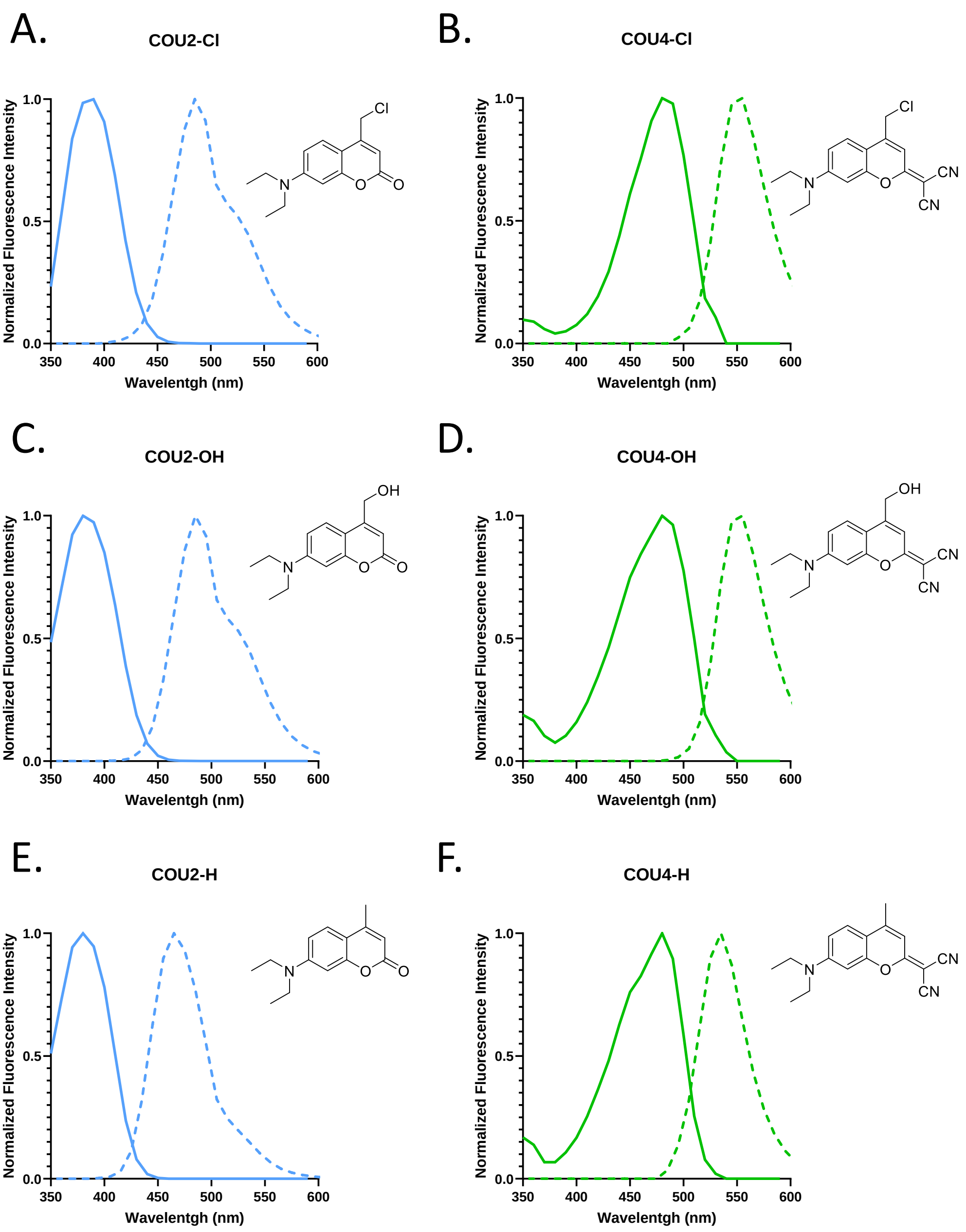

**Supplementary Figure 1**. Absorption and emission spectra of fluorescent substrates. Normalized fluorescent intensity of absorption (solid lines, $\lambda_{em(A, C)}$ = 495nm, $\lambda_{em(E)}$ = 465nm, $\lambda_{em(B, D)}$ = 545nm, $\lambda_{em(F)}$ = 535nm) and emission (dashed lines $\lambda_{exc(A, C)}$ = 390nm, $\lambda_{exc(E)}$ = 380nm, $\lambda_{exc(B, D, F)}$ = 480nm) spectra of 10 μM **COU2-Cl** (A), **COU2-OH** (C), **COU2-H** (E), **COU4-Cl** (B), **COU4-OH** (D), **COU4-H** (F). All spectra measured in phosphate buffer/DMSO mixture (90:10, v/v), pH 8.0.

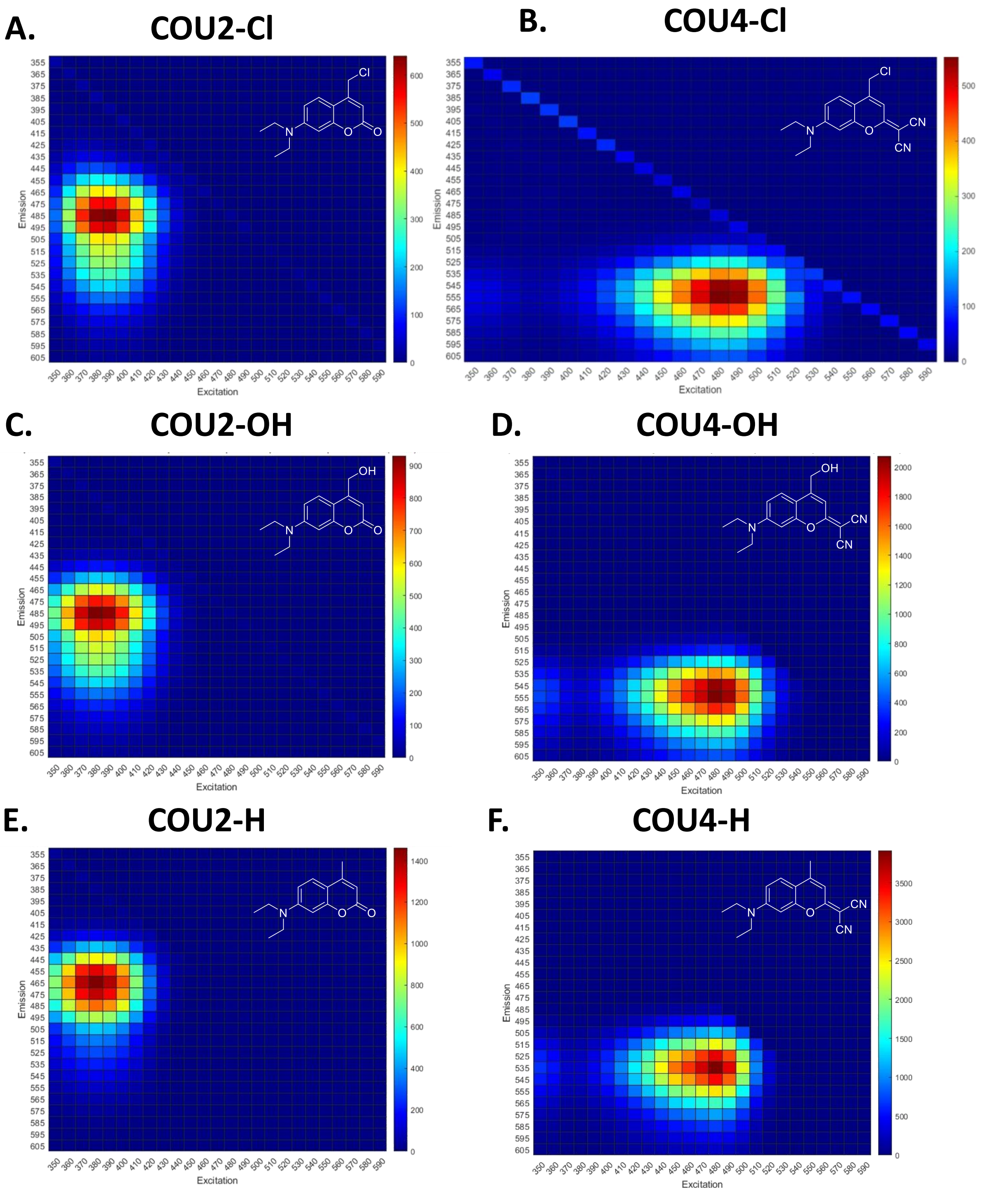

**Supplementary Figure 2.** 3D Excitation/Emission Spectra of **COU2-Cl** (A), **COU2-OH** (C), **COU2-H** (E), **COU4-Cl** (B), **COU4-OH** (D), **COU4-H** (F) at 10 µM in Phosphate Buffer 50 mM pH 8 10%DMSO (v/v). Large spectre (up) and zoom-in spectra (down). Fluorescence intensity is represented by a color scale from blue (low intensity) to red (high intensity).

**Specific activities of wild type HLDs toward COU2-Cl and COU4-Cl**

**Supplementary Table 1.** Specific activities of wild type HLDs toward COU2-Cl and COU4-Cl

| Enzyme | Specific activity (μmol $s^{-1}$ $mg^{-1}$) | |
|---|---|---|
| | COU2-Cl | COU4-Cl |
| DspoA | $(3.82 \pm 0.25) \times 10^{-1}$ | $(7.28 \pm 0.67) \times 10^{-1}$ |
| DphxA | $(1.78 \pm 0.13) \times 10^{-1}$ | $(1.31 \pm 0.14) \times 10^{-1}$ |
| LinB | $(1.73 \pm 0.21) \times 10^{-1}$ | $(5.54 \pm 1.11) \times 10^{-3}$ |
| DmmA | $(1.07 \pm 0.11) \times 10^{-1}$ | $(3.46 \pm 0.52) \times 10^{-2}$ |
| DexA | $(7.67 \pm 1.20) \times 10^{-2}$ | $(3.64 \pm 0.16) \times 10^{-3}$ |
| DadB | $(2.94 \pm 0.67) \times 10^{-2}$ | $(2.58 \pm 0.35) \times 10^{-2}$ |
| DprxA | $(1.33 \pm 0.01) \times 10^{-2}$ | $(1.84 \pm 0.39) \times 10^{-2}$ |
| DhxA | $(8.89 \pm 1.92) \times 10^{-4}$ | $(1.15 \pm 0.44) \times 10^{-3}$ |
| DbjA | $(7.62 \pm 1.65) \times 10^{-4}$ | $(1.41 \pm 0.20) \times 10^{-4}$ |
| DpcA | $(3.81 \pm 1.19) \times 10^{-5}$ | $(2.17 \pm 0.90) \times 10^{-3}$ |
| DhlA | $(1.75 \pm 0.07) \times 10^{-5}$ | $(7.45 \pm 1.66) \times 10^{-5}$ |
| DeaA | $(5.03 \pm 0.52) \times 10^{-6}$ | $(2.89 \pm 1.02) \times 10^{-5}$ |
| DbeA | $(2.88 \pm 1.10) \times 10^{-6}$ | $(2.67 \pm 1.52) \times 10^{-5}$ |
| DhaA | $(6.05 \pm 0.95) \times 10^{-7}$ | $(2.50 \pm 1.01) \times 10^{-5}$ |
| DmxA | n.d. | $(4.15 \pm 2.05) \times 10^{-6}$ |

n.d. not detected

**HPLC-FLD chromatograms of COU2-Cl and COU4-Cl digested by haloalkane dehalogenases**

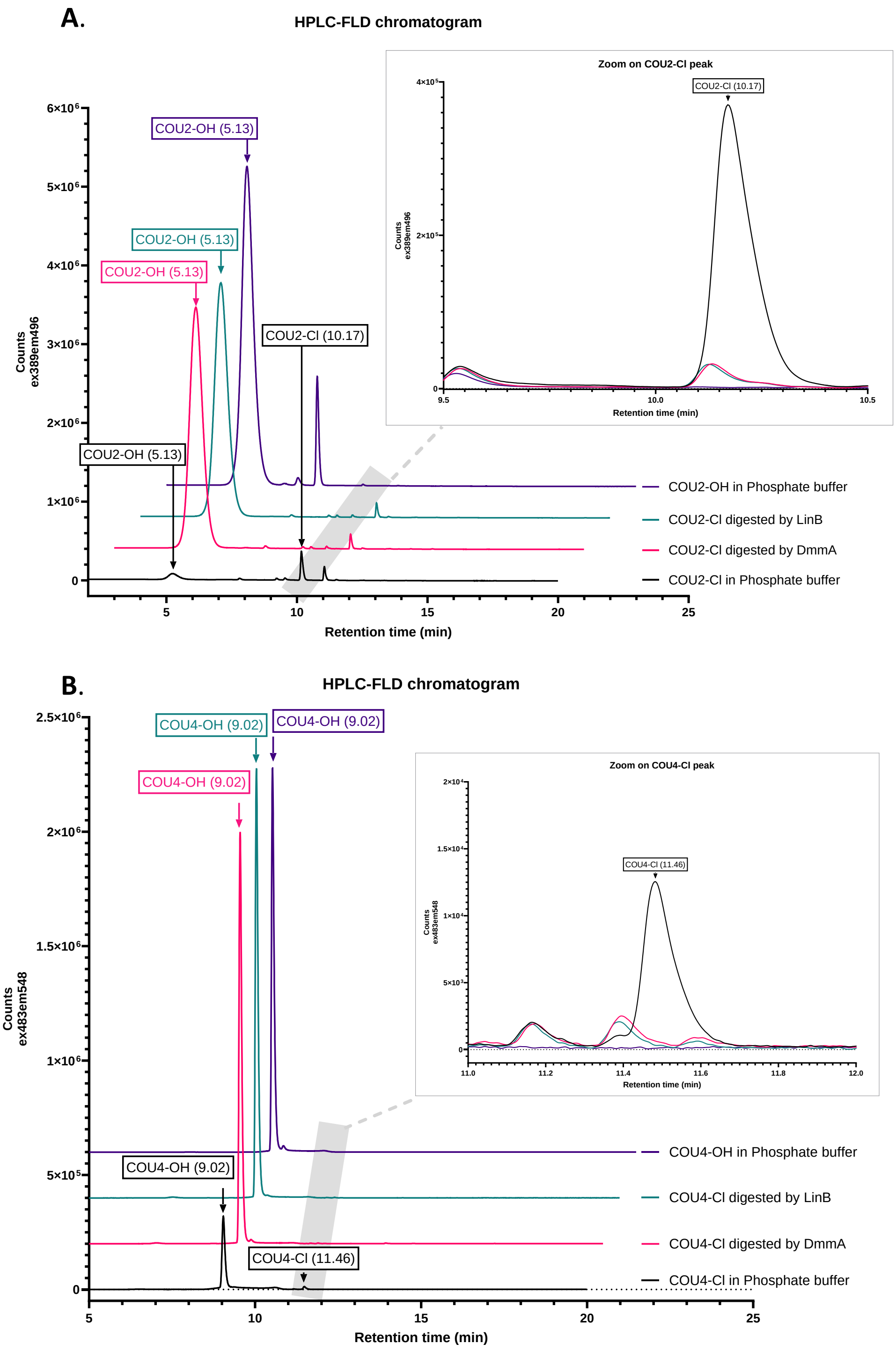

**Supplementary Figure 3.** HPLC-FLD chromatogram of 8 µM COU2-Cl (A) or COU4-Cl (B) after digestion by 10 µg.mL$^{-1}$ of LinB (green) or DmmA (pink), compared to control sample in phosphate buffer (black) and 8 µM of **COU2-OH** (A) or **COU4-OH** (B) (purple). $\lambda_{excA}$ = 389

nm, $\lambda_{emA}$ = 496 nm, $\lambda_{excB}$ = 483 nm, $\lambda_{emB}$ = 548 nm. Labels contain the name of identified molecules and the corresponding retention time. To allow for better results display, injection peaks are not shown, and each chromatogram is delayed on (A) by +1 min on the X axis and +400,000 counts on the Y axis, on (B) by +0.5 min on the X axis and +200,000 counts on the Y axis compared to the precedent. A spontaneous hydrolysis of the substrate is observed for **COU2-Cl** and **COU4-Cl** in phosphate buffer after incubation leading to the observation of **COU2-OH** and **COU4-OH** in these samples where the OH product formed amount to 0.5% and 5% respectively.

**Kinetic of dehalogenase activity in whole cell by fluorescence detection using COU2-Cl or COU4-Cl as substrate.**

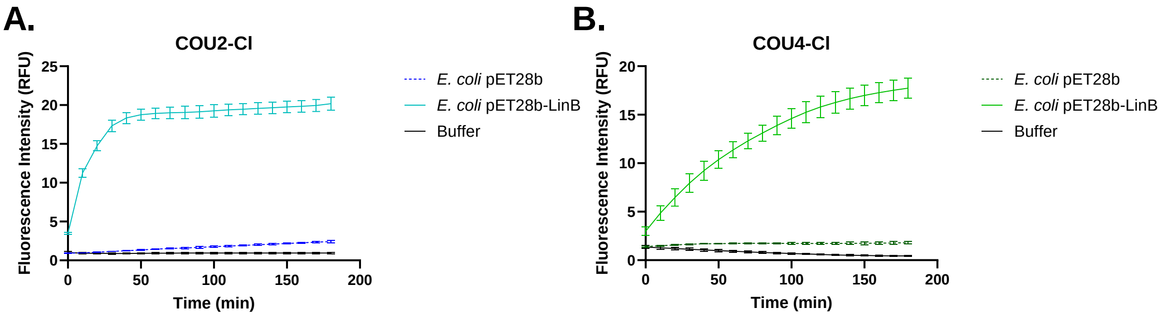

**Supplementary Figure 4.** Kinetic of dehalogenase activity in whole cell by fluorescence detection using **COU2-Cl** (A) or **COU4-Cl** (B) as substrate. Measurements were performed following the formation of **COU2-OH** (A) or **COU4-OH** (B) product by fluorescence at 30°C of *E. coli* cells with 8 µM **COU2-Cl** (A) or **COU4-Cl** (B), 10 % (v/v) DMSO in phosphate buffer at pH 8. n = 3 replicates for each condition. Fluorescence intensity measured in RFU, $\lambda_{exc(A)}$ = 389 nm, $\lambda_{em(A)}$ = 496 nm, $\lambda_{exc(B)}$ = 483 nm, $\lambda_{em(B)}$ = 548 nm.

#### Supplementary material: probe synthesis

*General information*

Reagents and solvents were supplied by Aldrich, Alfa Aesar and TCI Chemicals. NMR spectra were recorded on a Bruker 400 ($^{1}$H: 400 MHz; $^{13}$C: 100 MHz) spectrometers at 298 K, using $CDCl_3$ as solvent. The chemical shifts (δ, ppm) are referenced to the residual solvent peak and coupling constants (*J*) are reported in the standard fashion. The following abbreviations are used to explain the multiplicities: s = singlet, d = doublet, t = triplet, dd = doublet of doublet, m = multiplet. NMR peak assignments were performed for each compound using classic 1D and 2D NMR (COSY, HSQC and HMBC). High-resolution mass spectra (HRMS) were recorded on a Finnigan Mat 95xL mass spectrometer using electrospray (ESI). Analytical thin-layer chromatography was run on silica gel Merck 60 D254 (0.25 mm) plates. Column chromatography was performed on Merck Si 60 silica gel (40–63 µm). 7-(Diethylamino)-4-(hydroxymethyl)-2*H*-chromen-2-one, **COU2-OH** [1] 4-(chloromethyl)-7-(diethylamino)-2*H*-chromen-2-one, **COU2-Cl** [1] 2-(7-(diethylamino)-4-methyl-2*H*-chromen-2-ylidene)malononitrile, **COU4-H** [2] 2-(7-(diethylamino)-4-(hydroxymethyl)-2*H*-chromen-2-ylidene)malononitrile, **COU4-OH** [3] were prepared according reported procedures.

**2-(4-(Chloromethyl)-7-(diethylamino)-2H-chromen-2-ylidene)malononitrile, COU4-Cl**

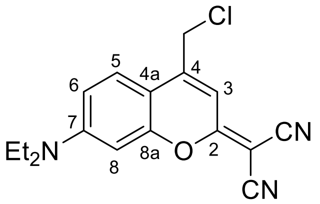

$PCl_5$ (35.3 mg, 0.169 mmol) was slowly added to a solution of **COU4-OH** (50 mg, 0.169 mmol) in $CH_2Cl_2$ (1 mL) at 0°C under argon. The reaction mixture was stirred at 0°C for 3h, and then $H_2O$ (10 mL) was carefully added to quench the reaction. The product was extracted with $CH_2Cl_2$ (3x20 mL), and the combine organic layers were washed with saturated $NaHCO_3$

(3x20 mL). The organic layer was dried over $Na_2SO_4$, filtered and the solvent was removed under reduced pressure. The crude product was purified by column chromatography (silica gel, Petroleum Ether:EtOAc, gradient 7:3 to 1:1) to give a pink-orange powder (33 mg, 62 %).

mp = degradation observed at 182°C.

$^1$H NMR (400 MHz, $CDCl_3$) δ 7.47 (d, *J* = 9.2 Hz, 1H, $H_5$), 6.76 (t, *J* = 0.8 Hz, 1H, $H_3$), 6.69 (dd, *J* = 9.2, 2.6 Hz, $H_6$), 6.59 (d, *J* = 2.6 Hz, $H_8$), 4.58 (d, *J* = 0.8 Hz, 2H, $CH_2Cl$), 3.46 (q, *J* = 7.1 Hz, 4H, 2 $CH_2$), 1.24 (t, *J* = 7.1, 6H, $2CH_3$).

$^{13}$C NMR (100 MHz, $CDCl_3$) δ 171.8 ($C_2$), 155.5 ($C_{8a}$), 151.9 ($C_7$), 146.4 ($C_4$), 125.7 ($C_5$), 114.5 (CN), 113.8 (CN), 110.8 ($C_6$), 108.6 ($C_3$), 107.4 ($C_{4a}$), 97.5 ($C_8$), 56.2 ($C_q$), 45.1 (2 $CH_2$), 41.1 ($CH_2Cl$), 12.6 (2 $CH_3$).

HR-MS (ESI, positive mode): m/z $[M+H]^+$ calculated for $C_{17}H_{17}ClN_3O$: 314.1055; found: 314.1054; m/z $[M+Na]^+$ calculated for $C_{17}H_{16}ClN_3NaO$: 336.0874; found: 336.0874.

*NMR Spectra*

**2-(4-(Chloromethyl)-7-(diethylamino)-2*H*-chromen-2-ylidene)malononitrile, COU4-Cl**

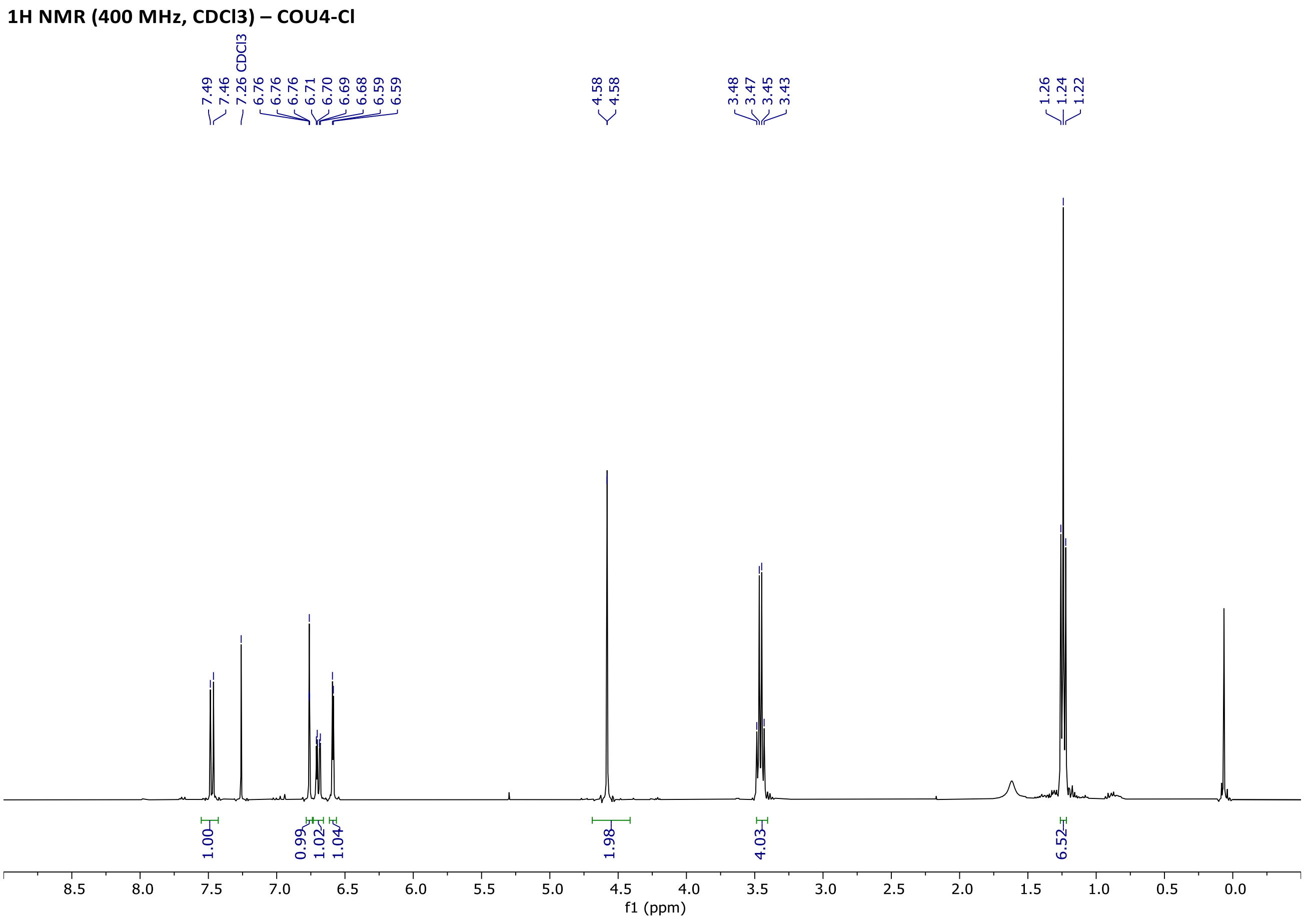

**13C NMR (100 MHz, CDCl3) – COU4-Cl**

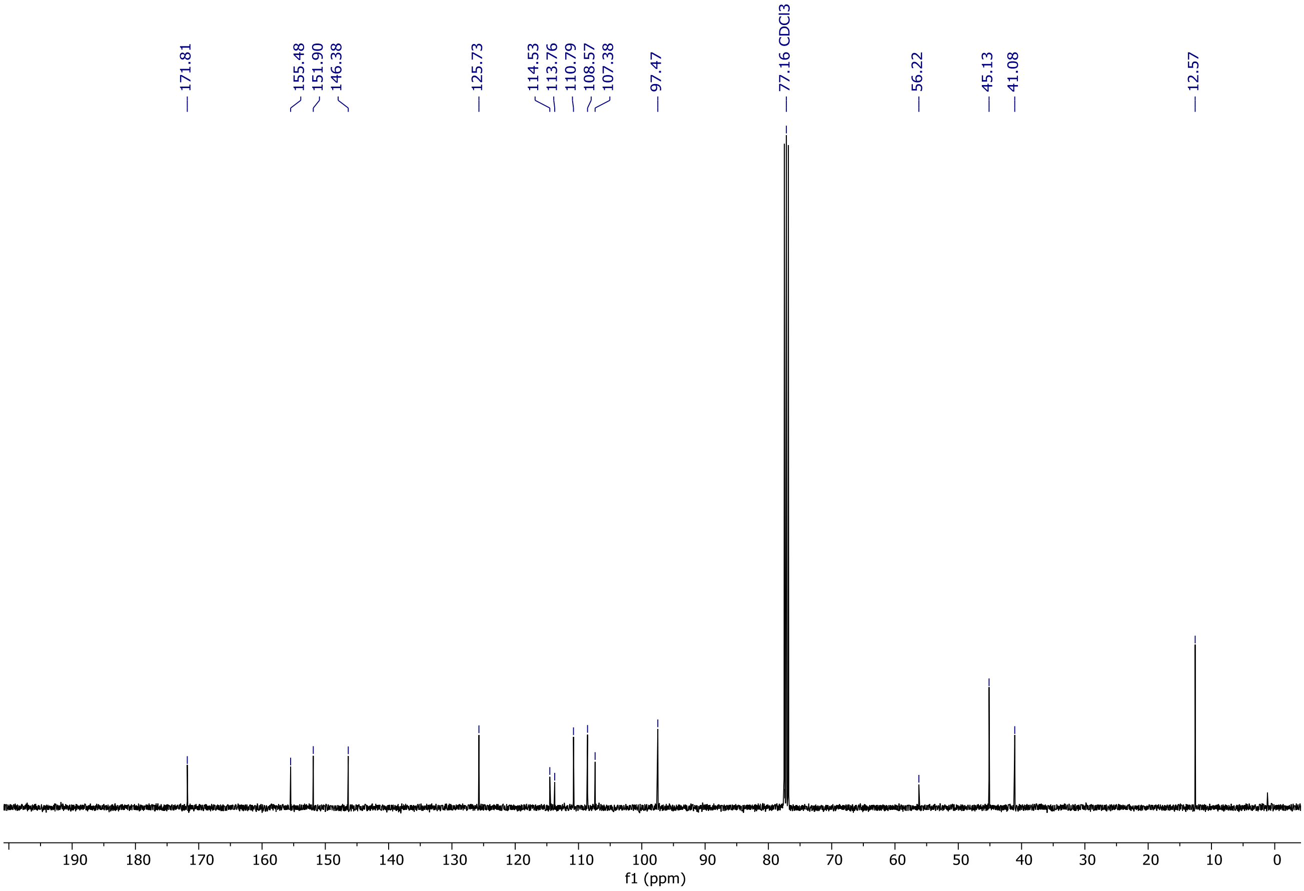

*HRMS Spectra*

**<u>2-(4-(Chloromethyl)-7-(diethylamino)-2*H*-chromen-2-ylidene)malononitrile, COU4-Cl</u>**

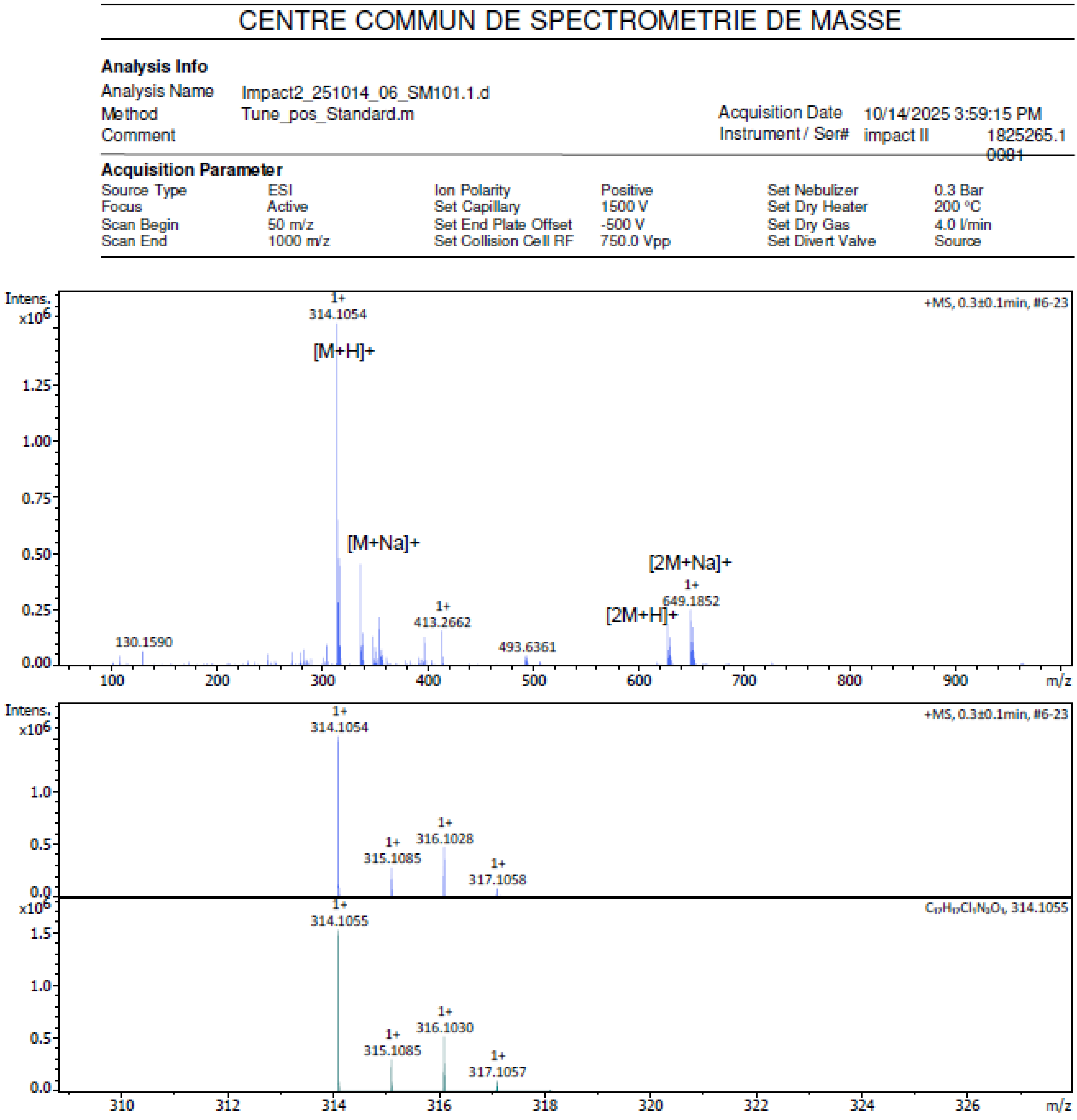

| Meas. m/z | Ion Formula | m/z | Sum Formula | err [ppm] | mSigma | Adduct | z |
|---|---|---|---|---|---|---|---|
| 314.1054 | C17H17ClN3O | 314.1055 | C17H16ClN3O | 0.1 | 14.9 | M+H | 1+ |
| 336.0874 | C17H16ClN3NaO | 336.0874 | C17H16ClN3O | 0.1 | 7.5 | M+Na | 1+ |
| 627.2033 | C34H33Cl2N6O2 | 627.2037 | C17H16ClN3O | 0.5 | 24.1 | 2M+H | 1+ |

#### Supplementary material: description of enzyme Cell Free Extracts (CFE) used in this study

**Supplementary Table 2.** Description of enzyme Cell Free Extracts (CFE) used in this study

| Enzyme | Origin from: | Reference |
|---|---|---|
| **DadB** | *Alcanivorax dieselolei* B-5, rec. from *E. coli* | [4] |
| **DbeA** | *Bradyrhizobium elkanii* USDA94, rec. from *E. coli* | [5] |
| **DbjA** | *Bradyrhizobium japonicum* USDA110, rec. from *E. coli* | [6] |
| **DhaA** | *Rhodococcus rhodochrous* NCIMB 13064, rec. from *E. coli* | [7] |
| **DhlA** | *Xanthobacter autotrophicus* GJ10, rec. from *E. coli* | [8] |
| **DmmA** | Marine microbial consortium, rec. from *E. coli* | [9] |
| **DmxA** | *Marinobacter* sp. ELB17, rec. from *E. coli* | [10] |
| **DpcA** | *Psychrobacter cryohalolentis* K5, rec. from *E. coli* | [11] |
| **LinB** | *Sphingobium japonicum* UT26, rec. from *E. coli* | [12] |
| **DeaA** | *Ensifer adhaerens*, rec. from *E. coli* | [13] |
| **DexA** | *Erythrobacter sp. NAP1*, rec. from *E. coli* | [13] |
| **DhxA** | *Hyphomona*s sp. BRH_c22, rec. from *E. coli* | [13] |
| **DphxA** | *Phaeobacter* sp. CECT 5382, rec. from *E. coli* | [13] |
| **DprxA** | *Pseudomonadota*, rec. from *E. coli* | [13] |
| **DspoA** | *Sulfitobacter pontiacus*, rec. from *E. coli* | [13] |

(6) Sato, Y.; Monincová, M.; Chaloupková, R.; Prokop, Z.; Ohtsubo, Y.; Minamisawa, K.; Tsuda, M.; Damborský, J.; Nagata, Y. Two Rhizobial Strains, Mesorhizobium Loti MAFF303099 and Bradyrhizobium Japonicum USDA110, Encode Haloalkane Dehalogenases

with Novel Structures and Substrate Specificities. *Appl. Environ. Microbiol.* **2005**, *71* (8), 4372–4379. https://doi.org/10.1128/AEM.71.8.4372-4379.2005

(12) Nagata, Y.; Hynková, K.; Damborský, J.; Takagi, M. Construction and Characterization of Histidine-Tagged Haloalkane Dehalogenase (LinB) of a New Substrate Class from a

Gamma-Hexachlorocyclohexane-Degrading Bacterium, Sphingomonas Paucimobilis UT26. *Protein Expr. Purif.* **1999**, *17* (2), 299–304. https://doi.org/10.1006/prep.1999.1143

(13) Vasina, M.; Vanacek, P.; Hon, J.; Kovar, D.; Faldynova, H.; Kunka, A.; Buryska, T.; Badenhorst, C. P. S.; Mazurenko, S.; Bednar, D.; Stavrakis, S.; Bornscheuer, U. T.; deMello, A.; Damborsky, J.; Prokop, Z. Advanced Database Mining of Efficient Haloalkane Dehalogenases by Sequence and Structure Bioinformatics and Microfluidics. *Chem Catal.* **2022**. https://doi.org/10.1016/j.checat.2022.09.011
